# Inference of high-resolution trajectories in single cell RNA-Seq data from RNA velocity

**DOI:** 10.1101/2020.09.30.321125

**Authors:** Ziqi Zhang, Xiuwei Zhang

## Abstract

Trajectory inference methods are used to infer cell developmental trajectories in a continuous biological process, for example, stem cell differentiation. Most of the current trajectory inference methods infer the developmental trajectories based on transcriptome similarity between cells, using single cell RNA-Sequencing (scRNA-Seq) data. These methods are often restricted to certain trajectory structures like linear structure or tree structure, and the directions of the trajectory can only be determined when the root cell is provided. On the other hand, RNA velocity inference method is shown to be a promising alternative in predicting short term cell developmental direction from the sequencing data. Here by we present `CellPath`, a single cell trajectory inference method that infers developmental trajectories by integrating RNA velocity information. `CellPath` is able to find multiple high-resolution cell developmental paths instead of a single backbone trajectory obtained from traditional trajectory inference methods, and it no longer constrains the trajectory structure to be of any specific topology. The direction information provided by RNA-velocity also allows `CellPath` to automatically detect the root cell and the direction of the dynamic process. We evaluate `CellPath` on both real and synthetic datasets, and show that `CellPath` finds more accurate and detailed trajectories compared to the state-of-the-art trajectory inference methods.

## Introduction

The availability of large scale single cell RNA-Sequencing (scRNA-Seq) data allows researchers to study the mechanisms of how cells change during a dynamic process, such as stem cell differentiation. One fundamental step in understanding the mechanisms is to reconstruct the trajectories of how cells change from one state to another. During recent years, various *trajectory inference* methods have been developed to perform this task^1–4^. These methods usually first learn the backbone structure of the trajectory, which can be linear, tree, cycle or other complex graph structure, and then each cell is mapped to the backbone and assigned a pseudotime.

Trajectory inference methods have led to significant biological discoveries, taking advantage of the large-scale, transcriptome-wide scRNA-Seq data^5–8^. However, due to that scRNA-Seq data captures only a snapshot of each cell in the cell population, although transcriptome similarity is used to find temporally neighboring cells, it is very hard to infer the direction of the trajectories using only the gene-expression profiles of cells. Most traditional trajectory inference methods ameliorate the loss of directional information by assuming developmental root cell is known as a prior, or using time-series data^7,9^. Moreover, the assumption that cells with similar gene-expression profiles should be sorted next to each other on the trajectory may not be true in real world scenario^10,11^.

Most traditional trajectory inference methods assume that the all cells in the dataset under analysis follow one trajectory structure, based on the backbone inferred. Methods were developed for specific topology of the backbone structure, including linear^12^, bifurcating^13^, tree-like^2^, and cycle structure^14^. Such constraints on the backbone topology confines these trajectory inference methods to be applicable to only a subset of real world dataset, and particularly those where there is only one starting point in the topology. In reality, a dataset can contain cells from multiple biological processes, which can correspond to a mixture of different topology types, or multiple trajectories with multiple root cells that cannot be covered by a defined backbone structure^15^. Even for cells within the same trajectory, cell sub-flows may exist, which creates multiple heterogeneous sub-trajectories^16^. Finally, even if the trajectory topology is fixed to be a certain type, some types including cycles and complex trees are particularly challenging for current methods^3^.

The recently developed RNA velocity methods^17,18^ can predict the gene-expression profile at the next time point for each cell by using the abundance of both nascent mRNA and mature mRNA. This information can potentially reveal “flows” of cell dynamics, which provides an alternative for resolving the loss of direction information in scRNA-Seq data. The packages `Velocyto`^17^ and `scVelo`^18^ provide visualizations on where the cells are moving to in 2-dimensional space. `Velocyto` plot an arrow following each cell pointing to its future state, and `scVelo` plot a streamline for a group of cells showing the dynamic trend in this group. However, none of these methods or tools output major cell trajectories in a dataset and the pseudotime of cells in each trajectory.

We hereby present `CellPath`, to bridge the gap between traditional trajectory inference methods and RNA velocity methods, as a method which outputs multiple high-resolution trajectories in a dataset using RNA velocity information. `CellPath` connects cells based on the future gene-expression profile of each cell predicted by RNA velocity, and identifies major paths in the dataset which correspond to main biological processes in the data. Taking advantage of the directional information of each single cell, `CellPath` overcomes certain problems of the traditional trajectory inference methods including the difficulty of automatically assigning trajectory directions and the restriction on the topology of the overall trajectory, and is applicable to datasets with any composition of biological processes.

The workflow of `CellPath` is shown in Fig. 1. `CellPath` takes as input the scRNA-Seq gene-expression matrix and RNA velocity matrix, which can be calculated from upstream RNA velocity inference methods such as `scVelo`^18^ and `velocyto`^17^. The various types of noise in scRNA-Seq data^19,20^ and noise in the estimated RNA velocity values^18^ pose challenges for the reconstruction of cell-level paths. It is common practice to construct “meta-cells”, which are clusters of cells, to reduce the effect of noise in each single cell^4,21,22^. `CellPath` follows the same route and starts with constructing meta-cells and performing regression model to obtain smoothed RNA velocity for each meta-cell (Fig. 1). The use of meta-cells can also reduce the computation complexity of the downstream trajectory detection. Then kNN graphs are constructed on the meta-cells, and we apply the Dijkstra shortest-path algorithm^23^ with constraints on the kNN graph to obtain a pool of possible trajectories within the dataset. Then, we design a greedy algorithm to select a small number of most likely trajectories within the pool, which give us the meta-cell level trajectories. To obtain cell-level trajectories and cell pseudotime, we develop an efficient named first-order pseudotime reconstruction method to assign order of single cells in each meta-cell separately and merge the orders together according to the meta-cell trajectories. `CellPath` is implemented as an open-source Python package (https://github.com/PeterZZQ/CellPath).

**Figure 1.**
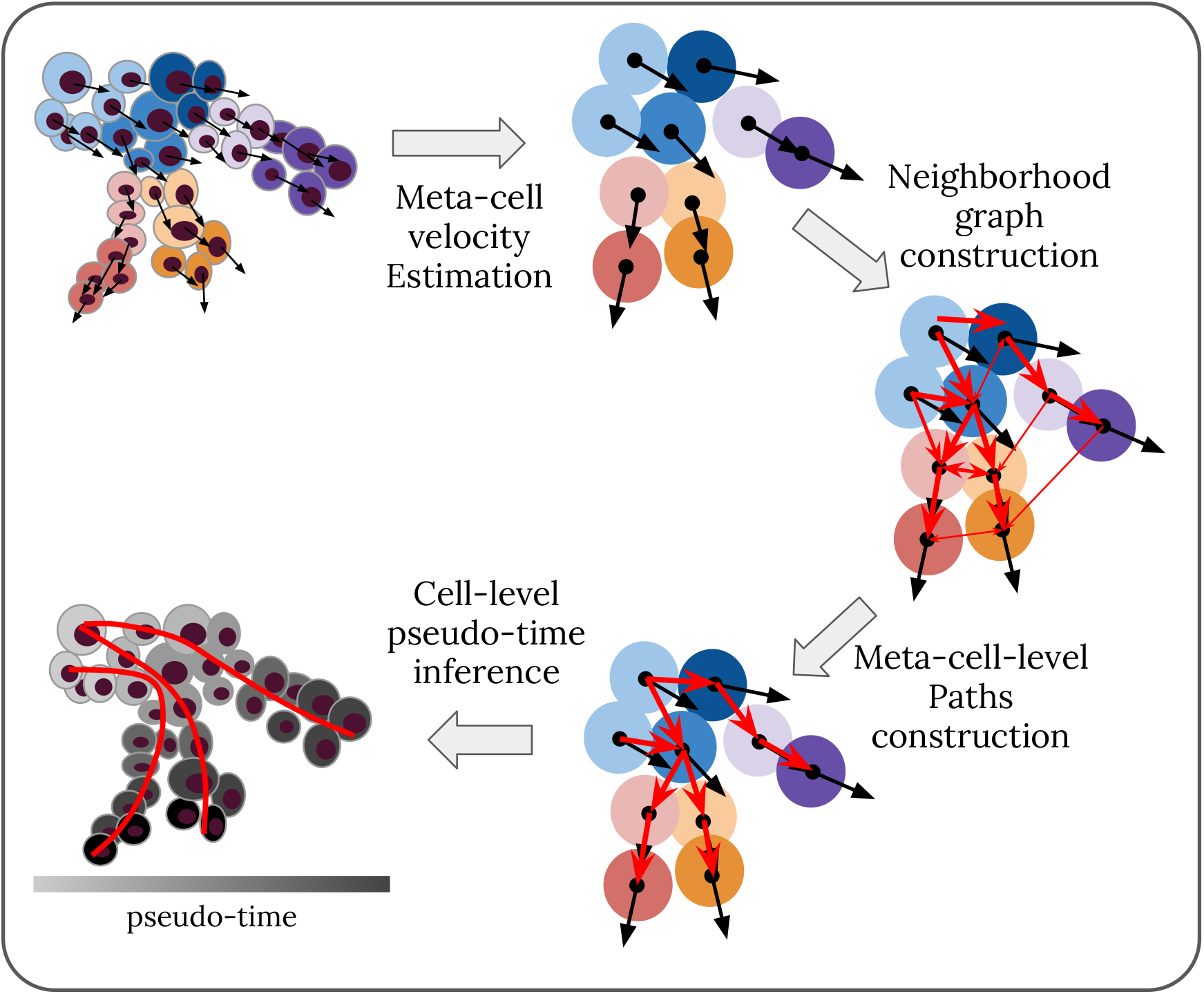
Workflow of `CellPath`. Step 1: `CellPath` constructs meta-cells and calculates their gene expression and RNA velocity profiles. Step 2: `CellPath` constructs a directed neighborhood graph on the constructed meta-cells. Step 3: `CellPath` uses a customized path-finding algorithm to find most probable meta-cell level trajectories on the neighborhood graph. Step 4: `CellPath` uses first-order pseudotime approximation algorithm to assign cell-level pseudotime.

We have tested `CellPath` on three real datasets and four different types of simulated datasets. The results verify the ability of `CellPath` in detecting subtle trajectories, which are often neglected in backbone-based methods, and in dealing with trajectories with complex structures including cycles.

## Results

We test `CellPath` on both real and synthetic datasets. We select real datasets with various levels of complexity in their trajectory structures. We apply `CellPath` on a dentate-gyrus dataset^15^ with 14 cell types and 2930 cells, a pancreatic endocrinogenesis dataset^24^ and a human forebrain dataset^17^ to analyse the performance of `CellPath`.

To be able to test `CellPath` with other trajectory topologies and to obtain quantitative measures on the performance of `CellPath`, we generate simulated data. We use two different tools to generate simulated data, `dyngen`^25^ and `VeloSim`^26^ (Methods) which can generate spliced counts, unspliced counts and the true RNA velocity with a given topology. These two simulators use very different principles for data simulation, where `dyngen` designs gene regulatory networks according to given trajectory backbone type, and generates mRNA counts with kinetic parameters controlled by the regulatory network; and `VeloSim` models the gene expression process using two states kinetic model, where the kinetic parameters are calculated based on cell identity vectors generated along the given backbone structure (Methods). The synthetic datasets include four different topology structures: a trifurcating structure, a multiple-batches bifurcating structure, a multiple-cycles structure, and a cycle-tree structure, where cells first go through a cell cycle process, and then differentiate into four different lineages. We also apply other state-of-the-art trajectory inference methods including `Slingshot, Vdpt` and `reCAT`^14^ for cell cycle process. The results on those datasets show that `CellPath` can detect high-resolution branching structure and is robust to various branching structure, even with highly complex branching structures, which are challenging cases for traditional trajectory inference methods.

### Results on Real data

#### CellPath captures major differentiation trajectories and subtle dynamic processes in dentate gyrus neurogenesis

To test the ability of `CellPath` in detecting complex lineage structure, we perform `CellPath` on a mouse dentate gyrus dataset^15^. The original paper where this dataset was published studied the dentate-gyrus neurogenesis process in developing and mature mouse dentate gyrus regions. The dataset has 2930 cells, and a UMAP^27^ visualization with cell types annotated is shown in Fig. 2a. The cells in this dataset are involved in multiple differentiation lineages which cannot be represented using a tree-like differentiation structure^15^. Therefore, most of the traditional trajectory inference methods which assume the trajectory has tree-like structures are not applicable to this dataset. `CellPath`, on the contrary, shows promising results on this dataset and detects both sub-paths in the cell dynamics in addition to all the mainstream differentiation lineages in the dataset.

**Figure 2.**
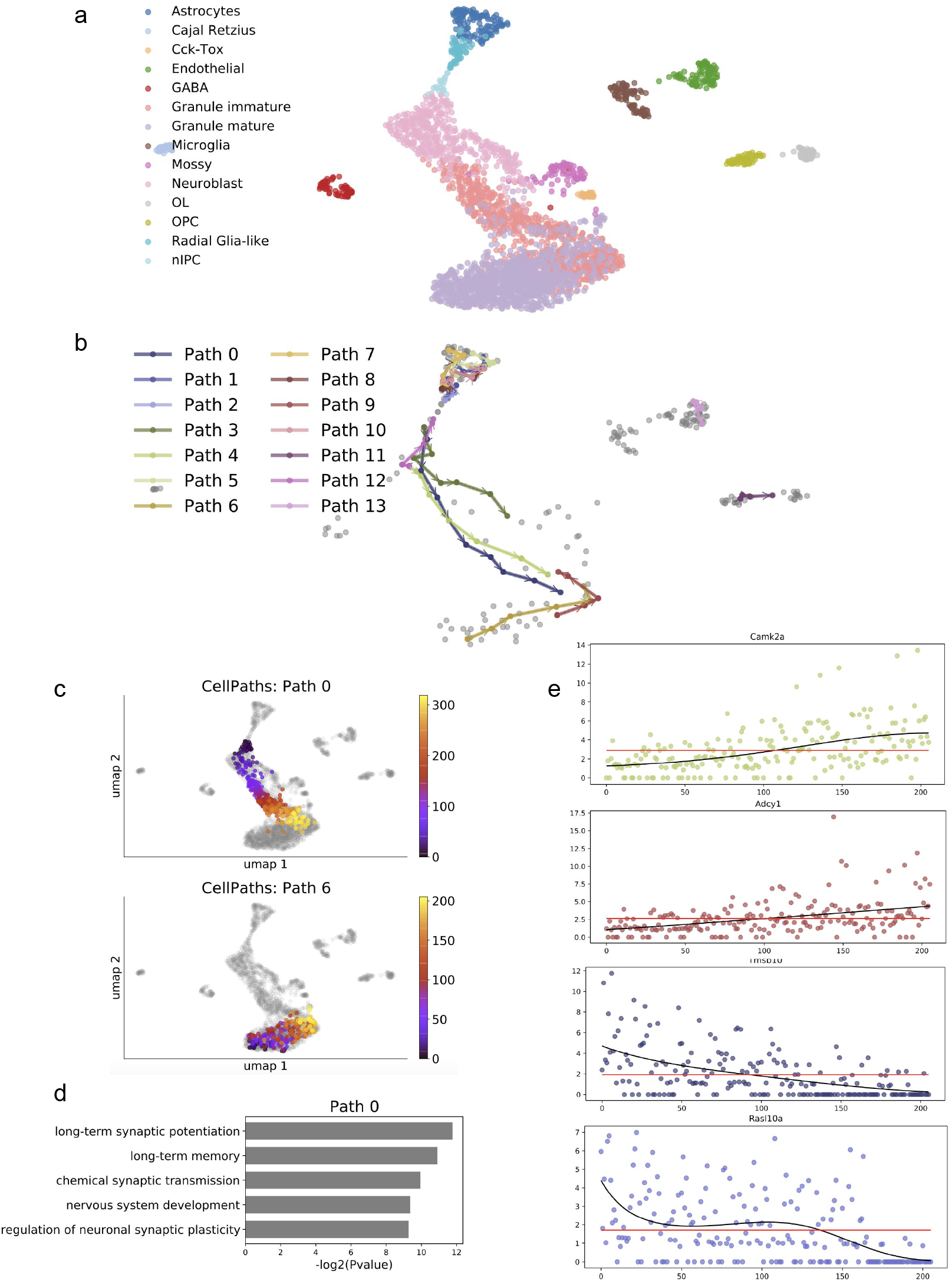
(a) Umap visualization of dentate-gyrus dataset with cell type annotated. (b) Meta-cell level paths inferred by `CellPath` on dentate-gyrus dataset. Gray dots corresponds to meta-cells. (c)Pseudotime inferred from `CellPath` on path 0 and path 6. The pseudotime is annotated by color. Yellow denotes smaller pseudotime, and purple denotes larger pseudotime. (d) Gene ontology analysis of DE genes on path 0. (e) The gene-expression level in terms of log(UMI counts) of DE genes *Camk2a*, *Adcy1*, *Tmsb10*, *Rasl10a* in cells sorted on Path 6.

In Fig. 2b, the top 14 paths that scores the highest according to `CellPath` greedy selection strategy (Methods) are shown at the meta-cell level. The algorithm infers multiple highly “time-coupled” trajectories, where the inferred trajectories consider both directional information of RNA velocity and transcriptome similarity, that follow the mainstream granule cells lineage, i.e. the differentiation path from neuronal intermediate progenitor cells (nIPCs), to neuralblast cells, immature granule cells and mature granule cells (Paths 0, 3 and 4). In addition, `CellPath` also detects paths corresponding to other small lineages: Radial Glia-like cells to Astrocytes (Paths 1, 2, 5, 7 and 8), Oligodendrocyte Precursor Cells (OPCs) to Myelinating Oligodendrocytes (OLs) (Path 11). Apart from the high-level lineages that correspond to distinct cell differentiation, `CellPath` also captures multiple small sub-flows of cells within the same cell-types, e.g. inferred trajectories within the mature Granule cell (Path 6) and Endothelial (Path 13).

We then calculate a pseudotime for each cell along the path it belongs to, using the “first-order approximation pseudotime assignment” method we propose (Methods). In Fig. 2c we show the cell pseudotime on Path 0 (the “nIPC–neuralblast– differentiation–immature granule–mature granule” cells differentiation path) and Path 6 (the mature granule internal path). Gene ontology (GO) analyses on the differentially expressed (DE) genes along each path are conducted to analyze the functionality of the inferred trajectories. With each inferred path, DE genes are detected by fitting a Generalized Additive Model (GAM) as a function of pseudotime to the gene-expression levels, and corresponding *p*-value is calculated using likelihood ratio test and corrected using false discovery rate (FDR) (Methods). Genes with p-value less than 0.05 are selected as DE genes. Gene ontology (GO) analysis is then conducted to summarize the function of genes within each path, with results shown in Fig. 2d. The result of the analysis further certifies the correctness of the inferred trajectories.

The 1st reconstructed trajectory in Fig. 2c (Path 0) corresponds to the main Granule generation process. The genes that change most abruptly with time are found to correspond well to the neuron morphogenesis, long-term synaptic potentiation(LTP) and neuron development, which shows an significant sign of neuron developmental process (Fig. 2d).

Interestingly, we found a path mostly inside the mature granule cells but end at the lower part of the immature granule cells (Path 6 shown in Fig. 2b). We identified genes that are differentially expressed (DE) on this path (Methods; the full list of DE genes are in Supplementary Table 1), and found multiple genes that may be relevant to the biological process along this path. *Camk2a* (also called the *α*-isoform of calcium/calmodulin-dependent protein kinase II) is known to be required for hippocampal long-term potentiation (LTP) and spatial learning and its deficiency can cause immature dentate gyrus, and other mental and psychiatric disorders^28–30^. The fact that we see its gene-expression increases within the mature granule cells may indicate the ongoing maturation of the granule cells or multiple subpopulation of the mature granule cells^31^. *Rasl10a* is reported to be exclusively expressed in the neuronal tissue and has a tumor suppressor potential^32^. The decrease of its expression level along Path 6 mostly represent the trend of the path going from the mature to immature granule cells, and the *Rasl10a* expresses highly only in the mature granule cells (Fig. 2e, Supplementary Fig. 1b). Neither of *Camk2a* or *Rasl10a* was discussed in the original paper of this dataset^15^ or in the paper where scVelo was applied to this dataset^18^. *Adcy1* may be involved in brain development and play a role in memory and learning (information from GeneCards^33^). *Adcy1* is known to be particularly highly expressed in granule cells in the brain^34^. *Tmsb10* is reported to be expressed in neural progentitors^35^ and this is in line with its expression level in this dataset (Supplementary Fig. 1a), but it is also expressed in some mature granule cells at the early stage of Path 6. The expression pattern of *Tmsb10* and the direction of Path 6 indicate that part of the mature granule cells may represent certain properties of the immature granule cells.

From the streamline visualization of RNA velocity in `scVelo` (Supplementary Fig. 1c, from paper^18^), one can see the trends of these paths but constructing the paths with `CellPath` allow us to extract the cells associated with each path and obtain the DE genes along each path.

#### CellPath captures cell-cycle and branching process in Pancreatic Endocrinogenesis

We further apply `CellPath` to a mouse pancreatic endocrinogenesis dataset^24^. The dataset profiles 3696 cells and includes endocrine cell differentiation process from ductal cells to four different endocrine cell sub-types, *α*, *β*, *δ* and *ε* endocrine cell, through Ngn3^low^ endocrine progenitor and Ngn3^high^ endocrine progenitor cell. The UMAP visualization of the dataset and corresponding cell-type annotation is shown in Fig. 3a. `CellPath` discovers multiple distinct lineages that correspond to *α*, *β* , *δ* endocrine cell genesis, with the meta-cell-level paths shown in Fig. 3b. Interestingly, we found a path where the *ε* cells turn into *α* cells (Path 7, Fig. 3b).

**Figure 3.**
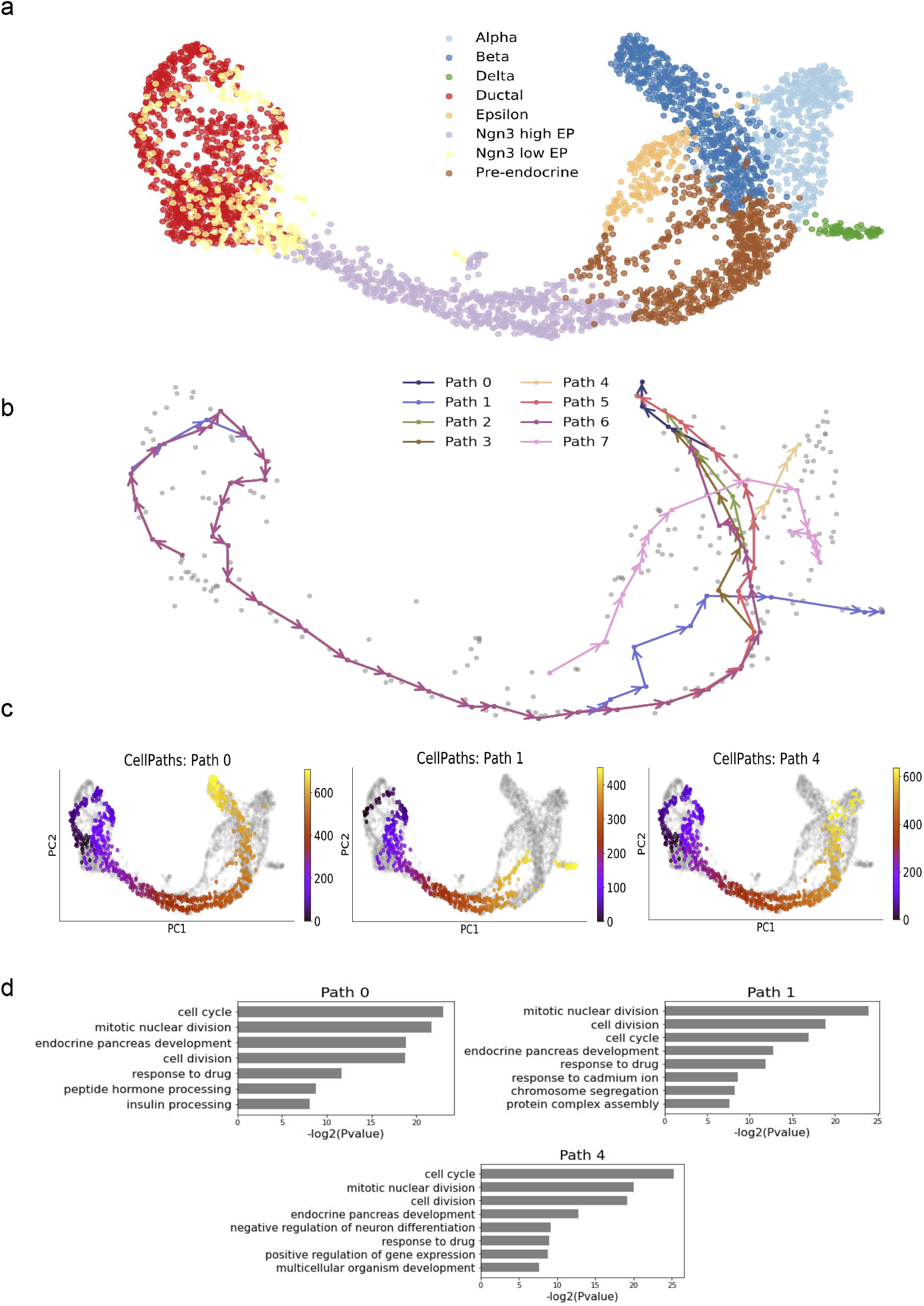
(a) Umap visualization of Pancreatic Endocrinogenesis dataset, with cell type annotated using different color. (b) Meta-cell level paths inferred by `CellPath` on Pancreatic Endocrinogenesis dataset. Gray dots corresponds to meta-cells. (c) Pseudotime inferred from `CellPath` on Paths 0, 1 and 6. The pseudotime is annotated by color. Yellow denotes smaller pseudotime, and purple denotes larger pseudotime. (d) Enriched GO terms of DE genes respectively on Path 0, 1 and 4.

DE gene analysis discovered multiple featured genes for different endocrine cell sub-types generation process, Within the insulin-producing *β* -cells generation trajectory, i.e. trajectory 1 in Fig. 3b, DE analysis discovers *Pcsk*2*, Ero*1*lb,Cpe* genes that function in insulin process. The strong time-correlationship of *Pax*4 is discovered in the somatostatin-producing *δ* -cells trajectory, trajectory 2 in Fig. 3b, which is known to have control over the endocrine cell type specification along with *Arx* and abundant in *δ* -cell lineage^36^. And within glucagon-producing *α*-cells generation path, trajectory 5, strong time-correlationship of *Arx* gene is discovered, which also correspond well to previous study^36^. Along with *Arx* and *Pax*4, a bunch of other highly time-coupled cell sub-type specific genes are discovered through `CellPath`, which allows for further study of cell sub-type specification mechanism. On the other hand, `CellPath` also discover a clear cell-cycle pattern on the left side of Fig. 3b, and multiple cell-cycle related genes are found through time-resolved DE analysis, such as *Ki f* 23*, Clspn, Aurkb, Spc*24, which further shows the high cell-level resolution of `CellPath` inferred trajectories.

#### CellPath finds multiple cell-flows in forebrain linear dataset

We further test `CellPath` on human forebrain glutamatergic neuron genesis dataset. The dataset profiles 1720 cells during the glutamatergic neuron differentiation process. Supplementary Fig. 2 shows a linear trajectory from Radial glia progenitor to fully differentiated neuron. `CellPath` is able to find multiple differentiation paths which are in line with the overall linear trajectory structure. All those paths correspond well to the glutamatergic neuron differentiation process while each path has their own developmental differences.

### Results on Simulated Data

#### Experiment design

To further evaluate `CellPath` performance compared to other state-of-the-art trajectories inference methods, we generate four synthetic datasets using two different simulators. To reduce possible bias of using only one simulator, we use simulators which use very distinct methodology. The first simulator, `dyngen`, generates the unspliced and spliced count matrix from customized gene regulatory network using Gillespie algorithm^37^; we use it to generate trifurcating structure and double-batch bifurcating structure. The second simulator is `VeloSim`^26^ (Methods). `VeloSim` generates spliced and unspliced counts using the two-state kinetic model, where a gene switches between the *off* and *on* states according to certain probabilities (Methods). Using `VeloSim`, we generate two datasets with complex trajectory structures. The first one is a “multi-cycle” structure where the trajectory traverses a cycle structure twice. The second one is a “cycle-tree” structure which consists of a cycle and a tree with three branching events. A biological example of this structure is where the cells first go through cell cycle and then start to differentiate into different cell types.

We compare our result with `Slingshot`, a method which shows state-of-the-art performance among all the other trajectory inference algorithms that utilize scRNA-seq data, according to the recent benchmark conducted by Saelens *et al*^3^ and Zhang *et al*^20^. We provide root cell information to `Slingshot` as a prior since it can not detect the root cell. In addition, we compare `CellPath` with trajectory inference methods that incorporate RNA-velocity information. Velocity diffusion pseudotime (Vdpt) is a trajectory inference method implemented in the `scVelo` package^18^ that is developed based on diffusion pseudotime^13^ and utilizes RNA-velocity for transition matrix construction and root-cell finding. The comparison shows that `CellPath` has the advantages of detecting high-resolution cell sub-flow like cell cycle within the branching trajectory, the ability of performing trajectories searching with complex structure with mixed topology, and the scalability for large datasets.

#### CellPath detects more correct branches in complex branching dataset

Bifurcation or multifurcation trajectory structures are often seen in cell differentiation processes^38,39^. It has been a challenge to accurately detect the branching point, and to distinguish between the branches which are relatively close. In order to test the performance of `CellPath` on common multifurcation trajectory structures, we generate two branching datasets using `dyngen`, one with a trifurcating structure, another with data from two batches, each with a tree structure.

We first compare our results with `Slingshot`^2^ on a simple trifurcating dataset. With simple datasets, `Slingshot` has a comparative performance with `CellPath`. However, `CellPath` detects the branching point more accurately. In Supplementary Fig. 3, `Slingshot` misclassifies cells around the branching area into wrong branches, while in Supplementary Fig. 3, `CellPath` correctly classifies cells into the branches that they belong to. `CellPath` can detect trajectories with high resolution mainly due to the incorporation of RNA velocity information. RNA velocity predicts a cell’s potential differentiation direction, and cells with similar expression data can possibly have distinct differentiation direction. The incorporation of RNA velocity serves as additional information to separate cells in the branching area with a high resolution.

We design the second simulated dataset to have more complex structure. We run the process of generating the trifurcating structure twice, and obtain two highly similar and overlaid datasets. The two datasets are close to each other in the gene-expression space but have a minor difference branching with regarding the branching position. We then simply merge these two datasets which creates a setting that is particularly difficult for the detection of the branching point. We perform both `CellPath` and `Slingshot` on this dataset. In Supplementary Fig. 4, `CellPath` detects all four sub-branches and accurate corresponding branching points, while as in Supplementary Fig. 4, `Slingshot` merges two adjacency branches together and only detects two branches. With dimension reduction methods that cannot separate the trajectories into distinct spatial position, high-level clustering algorithms tend to cluster cells from different but closely located trajectories together, which results in fewer branch detection. Current cluster-level trajectory inference methods produce a trajectory inference result more robust to the noise within the scRNA-seq data compared to cell-level trajectory inference methods, but its performance are severely affected by the upstream dimension reduction and clustering method. And `Slingshot`, being one of those methods, tends to predict fewer branches and ignore detailed cell differentiation information. `CellPath`, on the other hand, utilizes meta-cell methodology, where clusters of moderate sizes are constructed. This strategy increases the robustness towards the expression measurement and RNA velocity calculation noise while also reduce the possibility of misclassification.

We further quantify the reconstruction accuracy of `CellPath` and other trajectory inference methods using Kendall rank correlation coefficient between the ground truth pseudotime and the predicted pseudotime by each respective method. In the trifurcating dataset and two batches branching tree dataset, the coefficient is measured on each individual branches inferred by `Slingshot` and `CellPath`. The result shows that `CellPath` provides better reconstruction accuracy, especially in more complex structure(Fig. 4, Supplementary Fig. 3, Supplementary Fig. 4, Supplementary Fig. 5).

**Figure 4.**
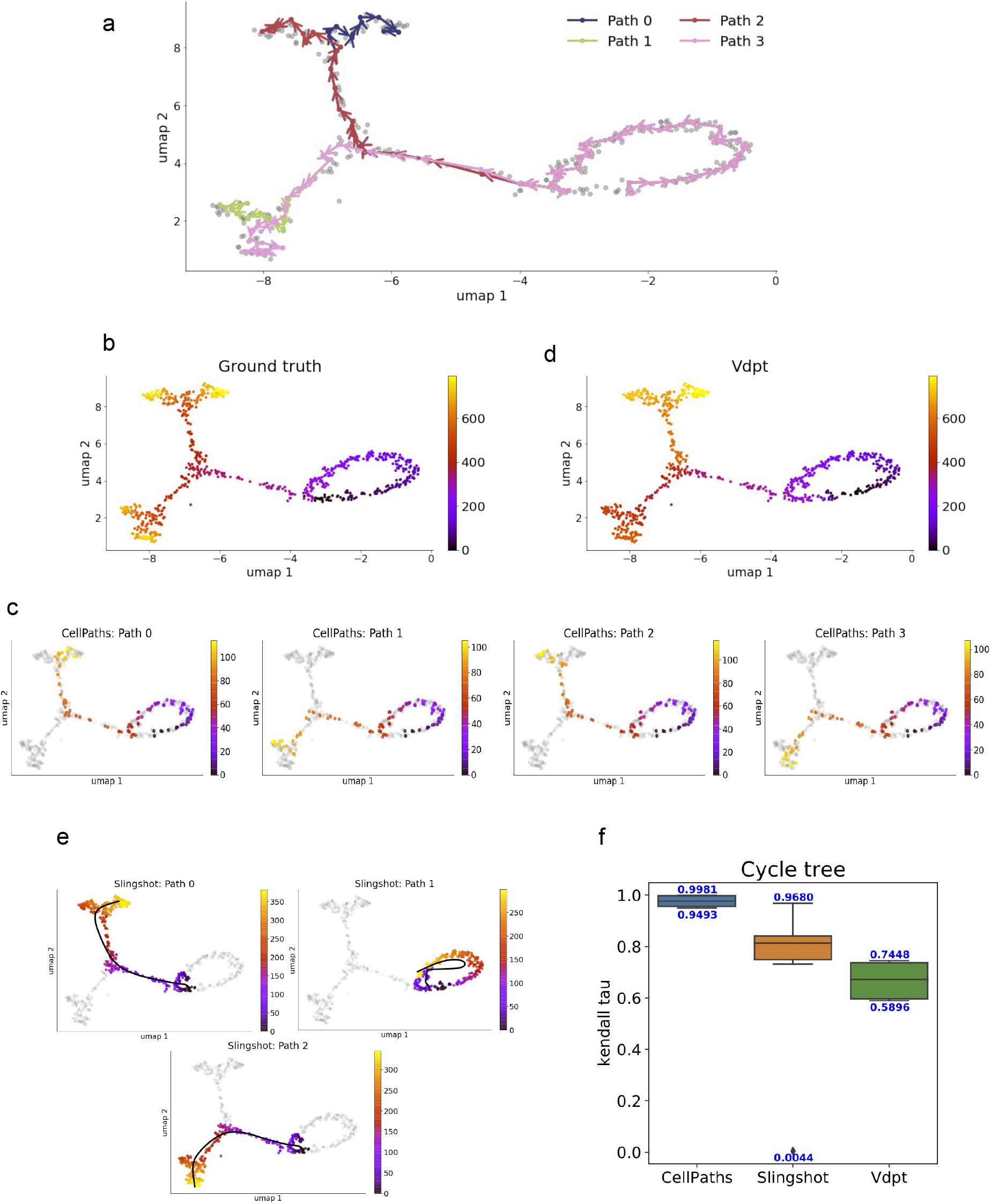
(a) Meta-cell-level paths generated by `CellPath` on the simulated cycle-tree dataset. The dataset is visualized using Umap. (b) Ground truth pseudotime annotation of the cycle-tree dataset. Yellow denotes smaller pseudotime, and purple denotes larger pseudotime.(c) Cell-level pseudotime of all four branches inferred by `CellPath`. The inferred pseudotime is annotated by different color. (d) The inferred pseudotime of `Vdpt. Vdpt` cannot differentiate cells in different branches. (e) The trajectory inferred by `Slingshot`. `Slingshot` infers two branches and one cell-cycle structure. The inferred principal curve and corresponding pseudotime of each branch is shown. (f) Boxplot of the Kendall rank correlation coefficient score of `CellPath`, `Slingshot` and `Vdpt`.

We use average entropy score (Methods), an entropy-based measurement we designed, to measure how well the inferred trajectory correspond to the ground truth differentiation path, namely the trajectory assignment accuracy.

We calculate the score on the two datasets above, where multiple real cell differentiation paths are generated are merged together, the average entropy score is shown in table 1. `CellPath` exhibit higher average entropy score, which shows that the trajectory assigned by `CellPath` is more similar to the real cell differentiation path compared to `Slingshot`, especially when the dataset has more complex branching structure.

**Table 1.** Trajectory assignment accuracy measured using average entropy score

|  | trifurcating | complex branching |
| --- | --- | --- |
| CellPath | 1.514 | 1.745 |
| Slingshot | 1.568 | 2.014 |

#### CellPath accurately infers cycle-structure in complex trajectory topology

In this section, we present the results of `CellPath` and other existing methods on two datasets which both contain cycle structures, one is referred to as “multi-cycle” and the other “cycle-tree”.

Detecting the cycle structures from a population of cells is shown to be challenging^3^. There are only a small number of methods which can detect the cycle structures and they tend to perform poorly^3^. The scenarios we generate here are more complex than a single cycle. In the “multi-cycle” structure we generate cells over a full cycle then continue to cycle and eventually form nearly two parallel cycles. We would like to test whether `CellPath` can find the cycle structure and further distinguish the two cycles. The “cycle-tree” structure is inspired by that some real world datasets can capture cells which are undergoing different biological processes, including cell cycle and cell differentiation. For example, the pancreatic data in^24^, cells first exit the cell cycle process and then enter the differentiation process. To generated a simulated dataset with similar scenarios, we use a topology where we have a complex tree with three branching events following a cycle structure (Fig. 4a).

`CellPath` successfully find all four branches and cell-cycle process (Fig. 4b-c). `Vdpt` by design infers only pseudotime of cells but not the trajectory structure. The pseudotimes it infers for the cells in the tree part are overall correct, but it did not distinguish the cells in the cycle part and those cells have very similar pseudotime. (Fig.4d). `Slingshot` finds three paths including the cell cycle path, but fails to infer the trajectory with only root cell provided.

Then, we perform `CellPath`, `Slingshot` and Vdpt on the “multi-cycle” dataset (Supplementary Fig. 5). As `CellPath` can accurately find multiple-cycle structure. On the other hand, `Slingshot` infer the differentiation process into a bifurcating structure without the RNA velocity information (given the true root cell). `Vdpt` and `reCAT` can accurately infer the differentiation direction, but it mixes cells from different cycles together and finds only one cycle (reflected in the relatively low Kendall rank correlation in Supplementary Fig. 5 which is further discussed in the next paragraph). In order to distinguish trajectory paths which are close, like the two cycles in the multi-cycle structure, the meta-cell size needs to be small, with a compromise on the noise reduction effect of larger meta-cells.

We then calculate the Kendall rank correlation to quantify the accuracy of cell pseudotime or ordering, for all three methods, `CellPath`, `Slingshot` and `Vdpt`. Multiple simulated datasets of each of the cycle tree and multiple cycles structures are generated using different random seeds, and the results of different algorithms are summarized using boxplots (Fig. 4, Supplementary Fig. 3, Supplementary Fig. 4, Supplementary Fig. 5). The performance of `CellPath` is again better than `Slingshot` and velocity diffusion pseudotime. Especially in the multiple-cycle dataset, without the direction annotation, `Slingshot` tend to infer the cycle structure as bifurcating structure. As a result, `Slingshot` provides results with both positive and negative correlations. `Vdpt`, on the other hand, incorporate velocity information, but still provide almost random results as it mix the cells in two cycles together. `CellPath`, on the contrary, still provide accurate inference results as it successfully differentiates cells in two difference cycles and correctly detects the differentiation starting point.

## Discussion

We have presented `CellPath`, a method to detect multiple high-resolution trajectories in scRNA-Seq datasets. Noise in the single-cell RNA-seq data has always been one major problem for trajectory inference methods. Cell-level approaches^5,13,40^, can detect branching point in a high resolution, but is extremely sensitive to measurement noise, while cluster-level methods^2,4,41,42^ find more comprehensive lineage structures that robust to noise at the expense of the loss accuracy in branching point detection. The construction of meta-cell is a method that lies in between, by creating meta-cells, we denoise the original expression and velocity matrix while still preserve the detailed structure of the dataset.

The shortest-path algorithm and greedy selection strategy allow for the discovery of trajectories in a fully automatic way. Since shortest-path algorithm finds almost all possible trajectories, the method does not have any assumption on the underlying backbone structure. Greedy selection strategy assumes that the true trajectories structure of the dataset should have low average weights and cover the cells in the dataset as possible. The assumption fit in the real circumstance and synthetic dataset extremely well, which also proves the correctness of the assumption.

## Methods

### Estimating RNA velocity for each gene in each cell

For a given set of cells, our method takes as input three matrices: the unspliced mRNA count matrix, the spliced mRNA count matrix, and the RNA velocity matrix. Each matrix is of dimention *M* by *N*, where *M* is the number of genes and *N* the number of cells.

The RNA velocity matrix can be calculated by an existing method, such as `scVelo`^18^ and `velocyto`^17^. In the results presented in this manuscript, the RNA velocity matrix is calculated using the dynamical model of `scVelo`^18^.

### Meta-cell construction

RNA velocity estimation at single cell level can be very noisy and even erroneous, given the noisy measurements of the count matrices especially the unspliced mRNA count matrix and the stringent assumptions on RNA velocity estimation. Even though current RNA velocity estimation methods take precautions to ameliorate the inaccuracy in estimation (e.g. `velocyto`^17^ and `scVelo`^18^ use k-nearest neighbor (kNN) graphs to denoise the measurement; `scVelo`^18^ relaxes the steady-state assumption of `velocyto`^17^ to dynamical model), using RNA velocity for trajectory inference can still suffer from the inaccuracy of upstream RNA velocity calculation. Here we propose to perform meta-cell construction as an denoising step prior to finding the trajectory paths

We assume the single cell gene-expression data that share strong similarities in the expression space are the noisy realizations of the underlying *meta-cell* gene-expression profile^21^. Meta-cells are constructed by clustering the single cells and deriving a profile for the meta-cell. Both k-means and Leiden clustering are implemented in `CellPath`, and k-means was used in the results we present. `CellPath` also provides the options of using both unspliced and spliced counts for clustering, or using only spliced counts for clustering. We have used both unspliced and spliced counts for the presented results.

For each meta-cell, its denoised gene-expression vector is calculated as the average of the gene-expression data of cells within the corresponding cluster. To obtain its smoothed RNA velocity measurement, we first construct a kernel regression model using the Gaussian radial basis function (RBF)^43^, *f* (**x**) = **v**, where the input 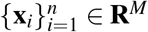 is the single cell gene-expression data (using spliced counts) and the output 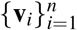 is the RNA velocity values, then use this function *f* (**x**) = **v** to calculate the meta-cell’s RNA velocity 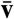 from its gene-expression profile 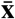.

In the process described above, *n* is the number of cells in the cluster corresponding to the meta-cell. This means that the smoothed RNA velocity measurement for a meta-cell is estimated based on the data within the cluster.

The kernel regression considers that the function lies within the reproducing kernel Hilbert space 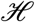 with projection 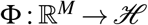. And the kernel function can be calculated using the projection 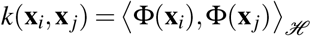.

We use Gaussian kernel which is one of the most widely used kernel smoothers for the regression model:

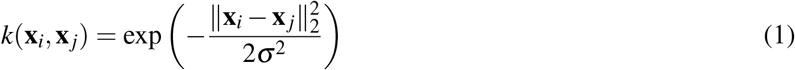

The final function is the linear combination of kernel functions

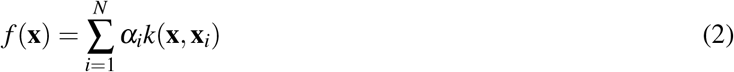

The coefficients *α* are calculated as *α* = (**K** + *δ* **I**)^−1^ **v**, derived from minimizing a MSE loss function including an L2-regularization term on *α*. The Kernel matrix **K** is simply calculated from the kernel function **K**_*i j*_ = *k*(**x**_*i*_, **x**_*j*_).

In our implementation, the kernel regression model *f* (**x**) = **v** is learned using the sklearn package in Python. The parameter *δ* which controls the regularization on *α* was set to be 1.

### Neighborhood graph construction

The cell differentiation mechanism can be modeled mathematically as a low dimensional manifold within a continuous high dimensional expression space^10,44^, which provide a strong theoretical support of manifold learning method in single-cell data analysis. Currently, manifold-learning-based methods^13,45,46^, for single-cell dataset construct neighborhood graph with different kinds of kernels to approximate the underlying manifold, which achieves promising results in single-cell dataset.

Construction of neighborhood groups are commonly used in single cell RNA-seq data analysis prior to graph-based clustering methods^4,47^. In existing work, the neighborhood graph construction process uses distance or similarity measurements of gene-expression profiles between cells and yields an weighted undirected graph. In our work, the RNA velocity information provides direction information on where each cell is going to next. To incorporate the direction information, we construct a weighted directed graph that penalize both the “direction difference” (detailed below) and transcriptome distance between every two cells. The graph construction process can be separated by two steps: *k*-nearest neighbor graph construction with selected *k*, and weight assignment to the edges in the kNN graph.

To calculate the direction penalty on an edge from cell *i* to cell *j*, we first define an angle *θ*. This is the angle between the direction from cell *i* to its future state defined by the RNA velocities of its genes, and the direction from cell *i* to cell *j*. We then define the direction penalty as *ℓ_θ_* (*i, j*) = 1 − cos(*θ*) where cos(*θ*) ∈ (0, 1], and

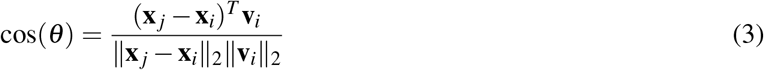

And distance penalty from cell *i* to cell *j* represents the transcriptome difference between the two cells in terms of spliced mRNA counts. It is calculated as

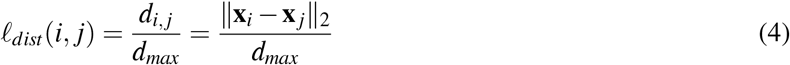

where 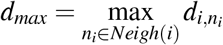, which is the largest distance from cell *i* to its neighbors. We have that *ℓ_dist_*(*i, j*) ∈ (0, 1].

Finally, the weight *e*(*i, j*) of an edge from cells *i* to *j* is calculated as following:

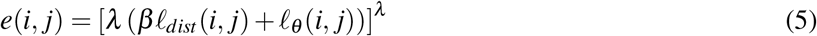

*β* and *λ* are hyper-parameters. *β* is used to adjust the relative contribution of the distance penalty and the direction penalty to the weight *e*(*i, j*), and *λ* is used to augment the difference between small and large weights. We set *λ* = 3 and *β* = 0.3 for all the results presented.

The graph construction pseudo-code is in the Supplementary Material.

### Detection of trajectory paths

Having constructed the weighted directed kNN graph on the meta-cells, we next detect trajectory paths in this graph which represent the cell dynamics in the dataset. We conduct two steps: first, we find a pool of candidate paths on the neighborhood graph, then we select the final paths using a greedy strategy as our reconstructed trajectories. The over aim is to find a small set of paths that cover as many vertices as possible.

Shortest-paths algorithms are suitable for weighted directed graphs to approximate the distance within the manifold between two vertices. However, shortest path algorithms can suffer from the noisy measurements, and the Floyd-Warshall algorithm which finds all-pairs shortest paths for a graph have *O*(*N*^3^) time complexity^48,49^. These problems are ameliorated through the following: 1) the use of meta-cells in the first step can increase robustness to noise; 2) instead of finding shortest paths between any two pairs for the pool, we limit the start vertices to be those with indegree at most 3 in the kNN graph, and then use the Dijkstra’s algorithm^23^ which finds shortest paths from a single start vertex to all other vertices. This practice accelerates the algorithm considerably and achieves comparable final results to those obtained using the Floyd-Warshall algorithm.

The pool of paths found with the procedure above can contain up to *N*^2^ paths. Next we would like to select a small number of paths which cover most of the vertices. We design a greedy path selection strategy which is conducted after initially removing some “bad paths”.

The paths that cover too few cells (the threshold varies with the total number of cells in the dataset), or have low average edge weights (with threshold 0.5) within the path are considered as “bad paths” and removed before he greedy selection. Shortest-path algorithm finds path between two nodes as long as those two nodes are connected. As a result, some directed shortest paths that connect two nodes but has large average edge weight usually have low time-coupling between neighboring nodes within the path and do not convey true biological causality relationship. The greedy algorithm picks paths iteratively and at each step it chooses the path with highest score, which is defined as for a path *p*: *S_p_* = *ζ* · *l_p_* + *l_u_*. *l_p_* is the length of path *p* in terms of the number of vertices in this path, and *l_u_* is the number of vertices which were not covered by any chosen path before choosing *p* but now are covered by path *p*. This means that the paths are selected based on both its own number of vertices and the number of vertices newly covered by this path. *ζ* is the parameter which finds a balance between *l_p_* and *l_u_*. With the greedy selection strategy, most meta-cells are covered by the first several paths.

The pseudo-code for greedy path selection algorithm is in the Supplementary Material.

### Assigning pseudotime to the cells on each trajectory path

Once we have the trajectory paths that cover the meta-cells, we proceed to assign pseudotime to the cells associated to the meta-cells on each path. Each meta-cell path can be considered as a linear trajectory structure for the cells covered by the meta-cells. Existing methods to assign cell-level pseudotime fall into two categories: principal-curve-based pseudotime assignment^2,12^ and random-walk-based pseudotime assignment^6,13,46^. However, these methods can not be readily used for our need and and they do not take advantage of the inferred meta-cell level paths, as root cell is the only information that is provided. Here we propose a *first order approximation* pseudotime assignment method, which is an efficient method with linear complexity.

After obtaining meta-cell paths, the relative order between meta-cells is known, and we need to assign orders for cells within each meta-cell. We can consider each predicted trajectory path as a smooth curve that passes through the “center” of each meta-cell on this path. The meta-cell center corresponds to the meta-cell gene expression 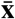 which is the denoised version of all the cells within the meta-cell. We denote the smooth curve by a function **f**(*t*) : ℝ → ℝ^*M*^, where *t* is the pseudotime and *M* is the number of genes. As **f**(*t*) passes through all the meta-cell centers, for any meta-cell *i*, there exists a point on the curve with **f**(*t_i_*) = **x**_*i*_, and the derivative of **f**(·) at *t_i_* is the RNA velocity **v**_*i*_ of the meta-cell **x**_*i*_. Applying first order Taylor expansion on **f**(*t*), we have

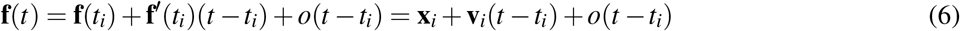

where *o*(*t* − *t_i_*) denotes the higher order derivative terms of **f**(*t*). When *t* is close to *t_i_*, we consider that *o*(*t* − *t_i_*) is small enough to be neglected, then we have **f**(*t*) ≈ **x**_*i*_ + **v**_*i*_(*t* − *t_i_*). This means that inside each meta-cell, the part of **f**(*t*) curve can be approximated by **g**(*t*) = **x**_*i*_ + **v**_*i*_(*t* − *t_i_*) which is a linear function.

Now for any cell *j* in the meta-cell (with center **x**_*i*_), to obtain its pseudotime, we project it to the linear function **g**(*t*) instead of the original function **f**(*t*) for which we do not have the analytical form.

Denoting the projected version of **x**_*j*_ by 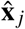, we have

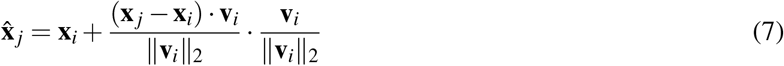

Note that the pseudotime we obtain is equal-spaced, meaning that we basically obtains the relative order between cells. Then for all cells in the same meta-cell, we simply compare their corresponding projected pseudotime 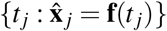 on **f**(*t*). It is obvious that the ordering of *t_j_* is the same as the ordering of the term 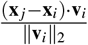 in Eq. 7.

Therefore, within each cluster, we calculate 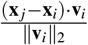, where **x**_*i*_ is the meta-cell expression, **v**_*i*_ is the velocity of the meta-cell, **x**_*j*_ is the true cells within the cluster, and then sort the result to obtain the ordering of cells in the meta-cell. We call this method to obtain cell ordering a *first order approximation* method.

In addition, principal curve and random walk based methods are also implemented in our `CellPath` package. We use mean first passage time^50^ as the pseudotime for the random walk based method.

### Differentially expressed gene detection and gene ontology analysis

A few methods were proposed to detect differentially expressed genes along a continuous trajectory. These methods generally test the significance of the expression level of a gene depending on a variable like pseudotime. Generalized linear models (GLM)^5^ and impulse models^51^ were used to model the dependency. Here we use a generalized additive model (GAM) which can model more patterns than GLMs (for example, where the expression of a gene first increases and then decreases).

The alternative hypothesis is that the gene-expression level *x* depends on pseudotime *t*. We assume the gene expression data follows a negative binomial distribution, and use spline function *f* () as the building block for the model, then we have

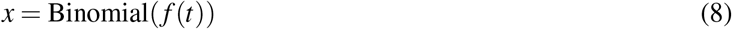

The null hypothesis is that the gene-expression level is irrelevant of the pseudotime, where we have

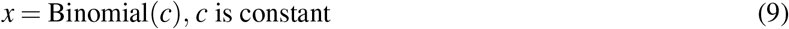

We test the two nested model in Eqs. 8 and 9 using likelihood ratio test. We test different genes one by one, and use false discovery rate (FDR) to correct the p-value for multiple testing and obtain adjusted p-values. We select deferentially expressed genes with p-value smaller than 0.05, and perform gene ontology (GO) enrichment analysis with `TopGO`.

### dyngen and VeloSim: generating simulated data

`VeloSim` is a procedure to simulate scRNA-Seq data including the amount of nascent RNAs and the true velocity^26^. Given a trajectory structure, `VeloSim` simulates time-series data of the expression levels of both the nascent and mature mRNAs. `VeloSim` follows the kinetic model^52^ and considers that a gene is either in an *on* state or in an *off* state. For every gene in every cell it generates the time course data of the gene’s expression in the cell, and samples the unspliced and spliced counts at a random time point to mimic the snapshot nature of scRNA-seq data.

### Evaluation metrics

We use two measures to evaluate the performance of trajectory reconstruction when ground truth information is available. On each trajectory path, we test whether the cells ordering we inferred is correct with Kendall rank correlation coefficient (Kendall’s Tau) measurement, and to test whether cells are assigned to the correct path, we use the *average entropy* score.

The *average entropy* scores are calculated for the data simulated by `dyngen`. At each simulation run, dyngen simulates the dynamics of one cell. When we repeat this process multiple times, we get multiple trajectories, which are almost identical to each other, if we keep all the parameters the same. (Key parameters include the backbone type, number of TF and target genes and kinetic parameters.) We show that `CellPath` can separate these trajectories, each theoretically corresponding to the developmental path of one cell.

The average entropy measurement is calculated as follows: for each inferred trajectory, we take the cells inferred to be on this trajectory, and group these cells according to their ground truth trajectory origin. Then we calculate the proportion of cells that belong to different ground truth trajectory, and obtain a discrete distribution. We then calculate the entropy of this distribution. That is, for inferred trajectory *j* ∈ **J**, denoting the cells that belong to simulation *i* by **S**_*j*_(*i*), then the proportion and entropy of this trajectory can be calculated as

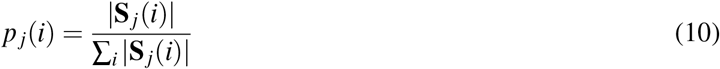

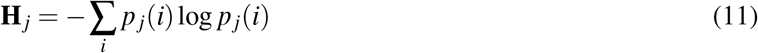

The final average entropy score is the average of entropy **H**_*j*_ over all *j* ∈ **J**.

Smaller average entropy correspond to better cell assignment to trajectories, with average entropy that equals to 0 corresponding to the ground truth assignment cell.

### Real datasets

We demonstrate the performance of `CellPath` using two previously published single-cell RNA-seq datasets, with RNA velocity calculated using the dynamical model in `scVelo`^18^.

#### Dentate Gyrus dataset

The original paper collected multiple dentate gyrus samples at different time points during mouse development^15^. The scRNA-seq process is performed using droplet-based approach and 10x Genomics Chromium Single Cell Kit V1. As in `scVelo`, we take the cells corresponding to the P12 and P35 time points from the original dataset. 2930 cells are incorporated that covers the full developmental process of granule cells from neuronal intermediate progenitor cells (nIPCs). The original data can be accessed through GSE95753.

#### Pancreatic Endocrinogenesis dataset

The dataset used to test `CellPath` is sampled from *E*15.5 of the original Pancreatic Endocrinogenesis dataset^24^. This dataset has 3696 cells and covers the whole lineage from Ductal cell through Endocrine progenitor cells and pre-endocrine cells to four different endocrine cell sub-types. The dataset is obtained through droplet-based approach and 10x Genomics Chromium. The original dataset can be accessed through GSE132188.

#### Human forebrain dataset

The dataset profiles 1720 using droplet-based scRNA-seq method, which incorporates cells spans from radial glia to mature glutamatergic neuron within glutamatergic neuronal lineage in developing human forebrain^17^. The original dataset is accessible with code SRP129388.

## Author Contributions

Z.Z. and X.Z. designed the algorithm, Z.Z. analysed the results. All authors wrote and reviewed the manuscript.

---

**Supplementary Figure 1.**
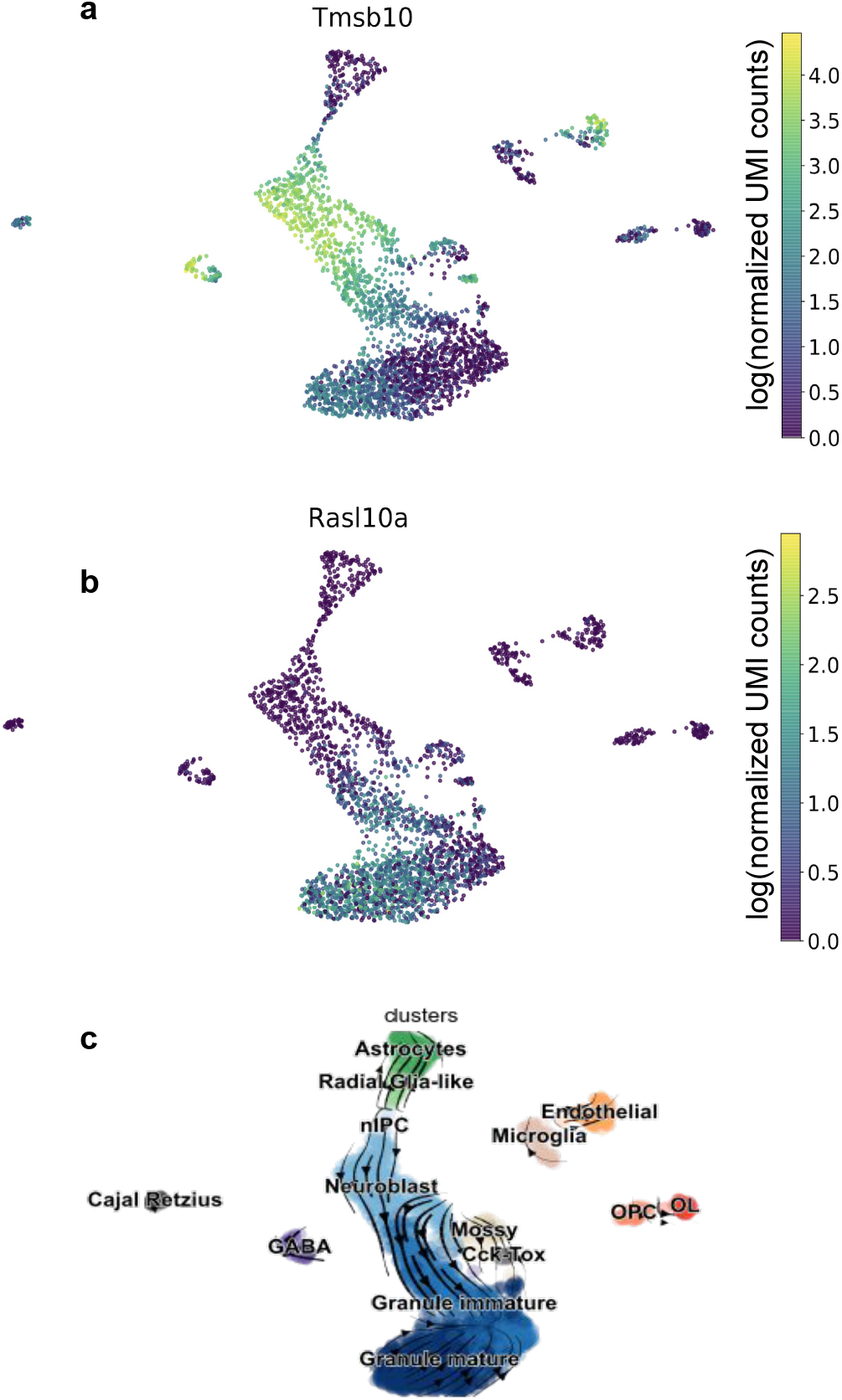
(a) Umap visualization of dentate-gyrus dataset colored by the expression level of *Tmsb10*. (b) Umap plot of dentate-gyrus dataset colored by the expression level of *Rasl10a*. (c) Streamline plot of dentate-gyrus dataset, streamlines correspond to the flow of the RNA velocity plotted by `scVelo`^18^.

**Supplementary Figure 2.**
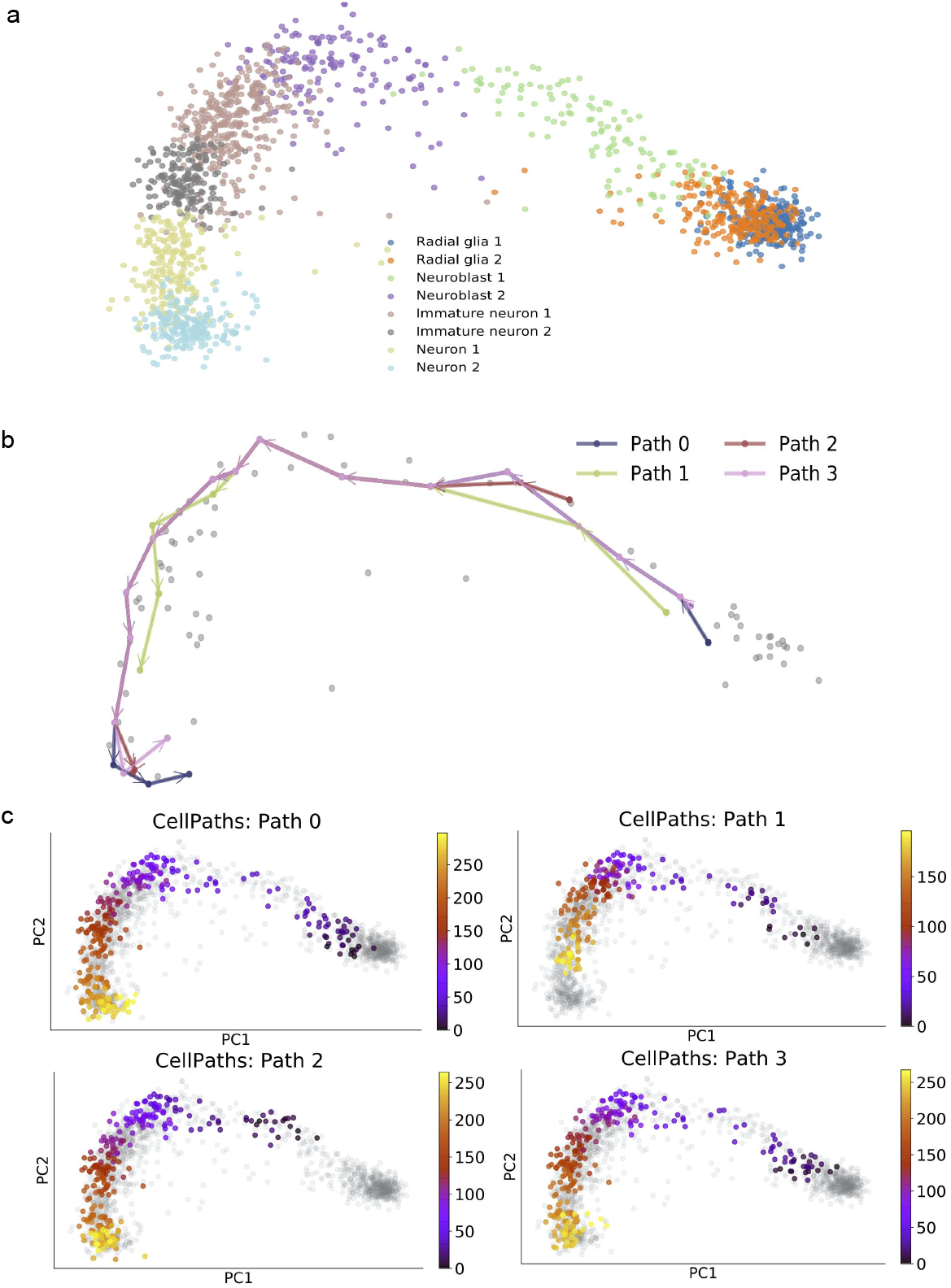
(a) PCA visualization of Human forebrain glutamatergic neuronal lineage dataset, with cell type annotated using different colors. The dataset shows a linear trajectory from radial glia to fully differentiated glutamatergic neuron. (b) Multiple meta-cell paths within the linear trajectory are discovered using `CellPath`. (c) Cell-level pseudotime of each meta-cell path is inferred using `CellPath`. The gradual change of cell states corresponds well to the differentiation process of human forebrain glutamatergic neuron.

**Supplementary Figure 3.**
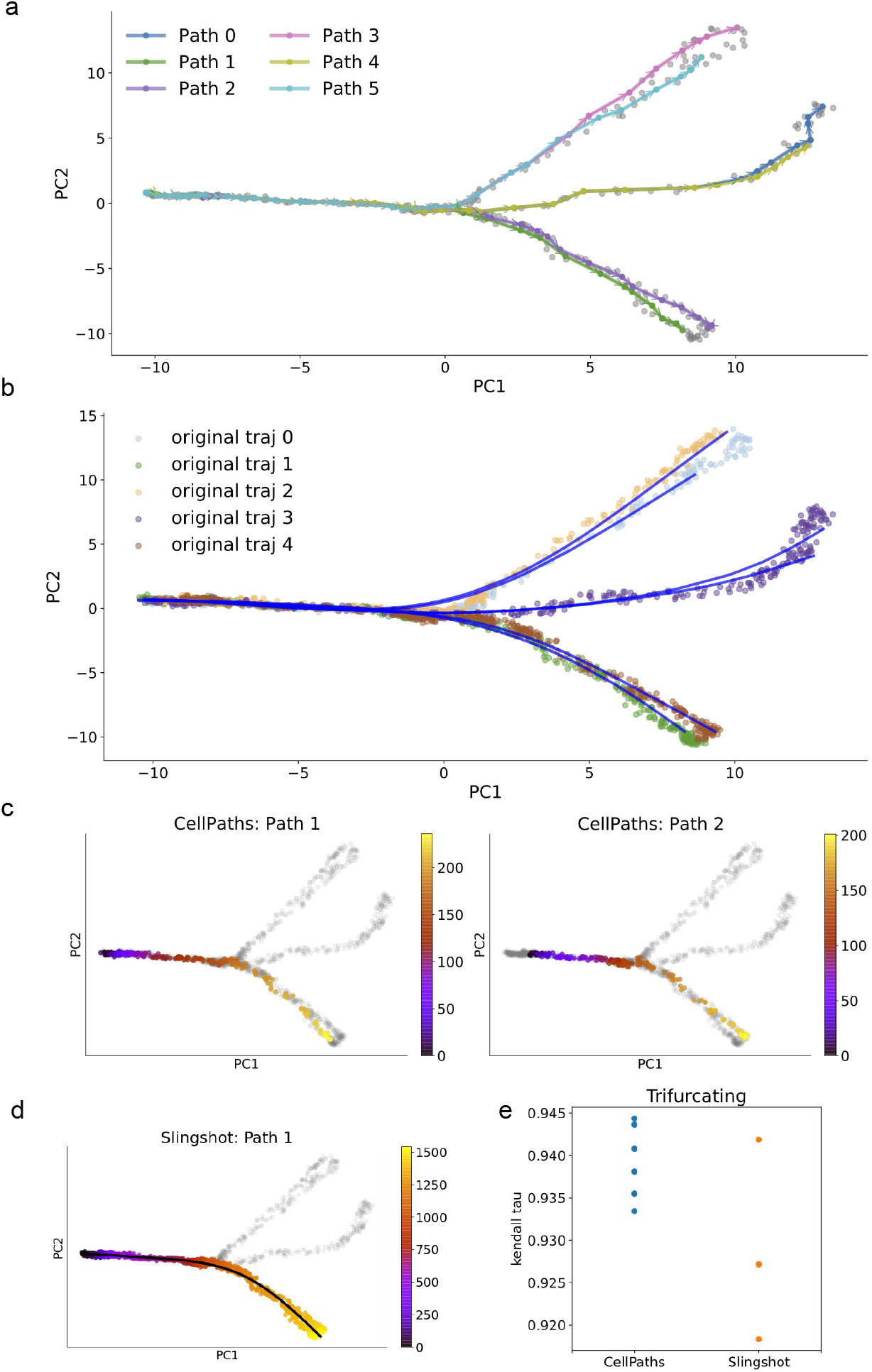
(a) The meta-cell level paths detected using `CellPath` on simulated trifurcating dataset. The dataset is visualized using PCA, and gray dots corresponds to different meta-cells. (b) Principal curve visualization of the trajectories inferred by `CellPath`, Dots correspond to true cells, and the cells that belong to different ground truth trajectories are colored differently. (c)(d). The pseudotime ordering of cells in one branch(colored from yellow to purple) inferred by `CellPath` and `Slingshot`. `CellPath` discovers two trajectories within the branch, while `Slingshot` only discovers one. The color(from yellow to purple) represent the ordering of cell in current trajectory, yellow denotes smaller pseudotime, and purple denotes larger pseudotime. Cells that do not belongs to current trajectory are colored gray. (e) Kendall rank correlation coefficient scores measured on the pseudotime inferred from `CellPath` and `Slingshot`. The score is calculated for each individual trajectory separately.

**Supplementary Figure 4.**
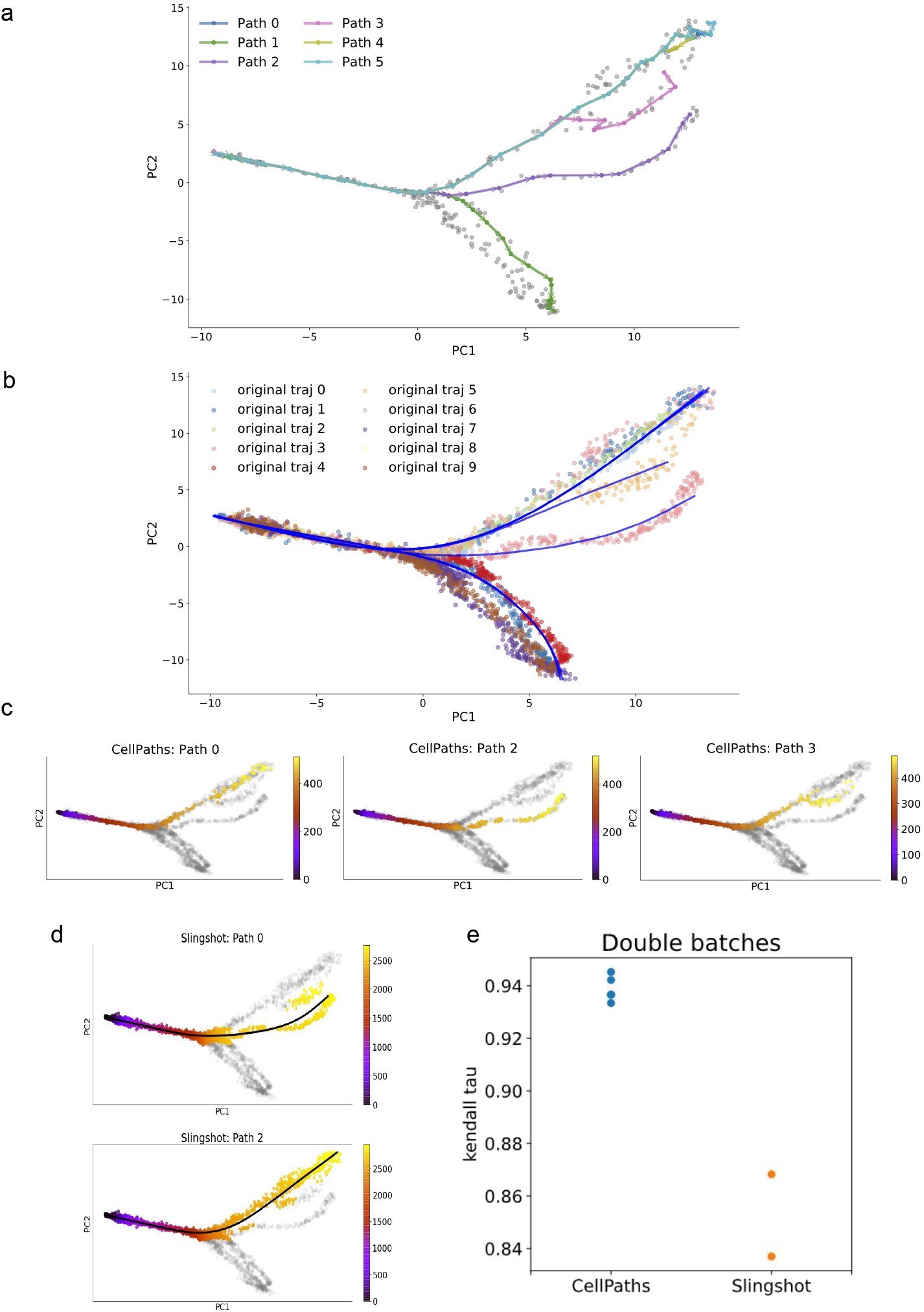
(a) The meta-cell level paths detected using `CellPath` on simulated complex branching dataset. The dataset is visualized using PCA, and gray dots corresponds to different meta-cells. (b) Principal curve visualization of the trajectories inferred by `CellPath`. Dots correspond to true cells, and the cells that belong to different ground truth trajectories are colored differently. (c)(d). The pseudotime ordering of cells in one branch(colored from yellow to purple) inferred by `CellPath` and `Slingshot`. `CellPath` discovers three trajectories within the branch, while `Slingshot` discovers two. The color(from yellow to purple) represent the ordering of cell in current trajectory, yellow denotes smaller pseudotime, and purple denotes larger pseudotime. Cells that do not belongs to current trajectory are colored gray. (e) Kendall rank correlation coefficient scores measured on the pseudotime inferred from `CellPath` and `Slingshot`. The score is calculated for each individual trajectory separately.

**Supplementary Figure 5.**
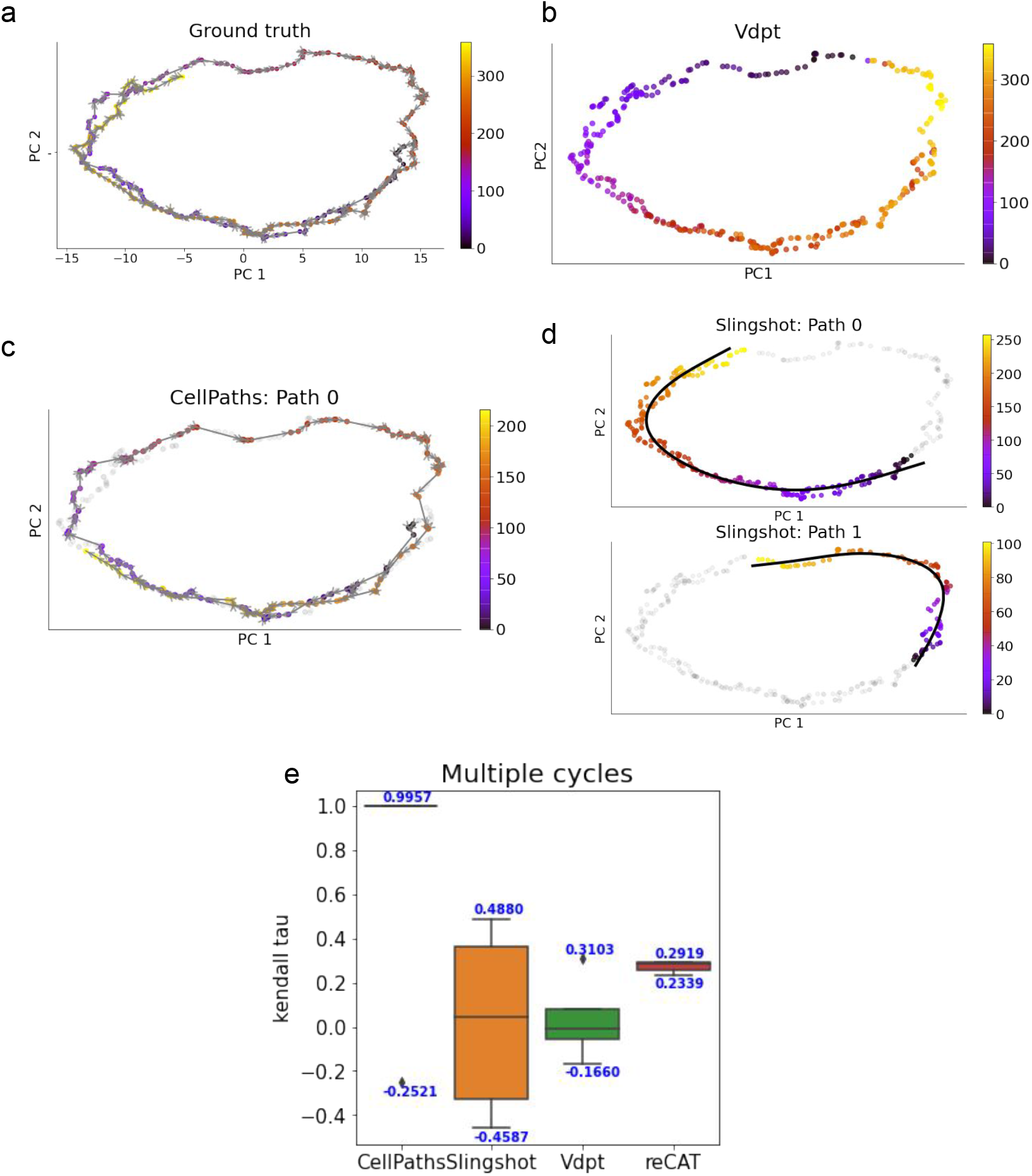
(a) PCA visualization of the multiple-cycle dataset. The cell color(from yellow to purple) represents ground truth pseudotime. Yellow denotes smaller pseudotime, and purple denotes larger pseudotime. The gray lines correspond to the ground truth trajectory backbone. (b) The pseudotime inferred by `Vdpt`. The pseudotime is annotated with different colors. (c) The pseudotime inferred by `CellPath`. Pseudotime is annotated with different colors, and adjacent cells are connected with gray line. The gray line shows that `CellPath` successfully infers two cycles within the dataset. (d) The trajectory inferred by `Slingshot`. `Slingshot` infers two branches. The inferred principal curve and corresponding pseudotime of each branch is shown. (e) Kendall rank correlation coefficient scores measured on the pseudotime inferred from `CellPath`, `Slingshot`, `Vdpt` and `reCAT`. Multiple datasets with the same structure are generated, and the results of multiple runs are visualized using boxplots.

## Supplementary Code

**Algorithm 1.**
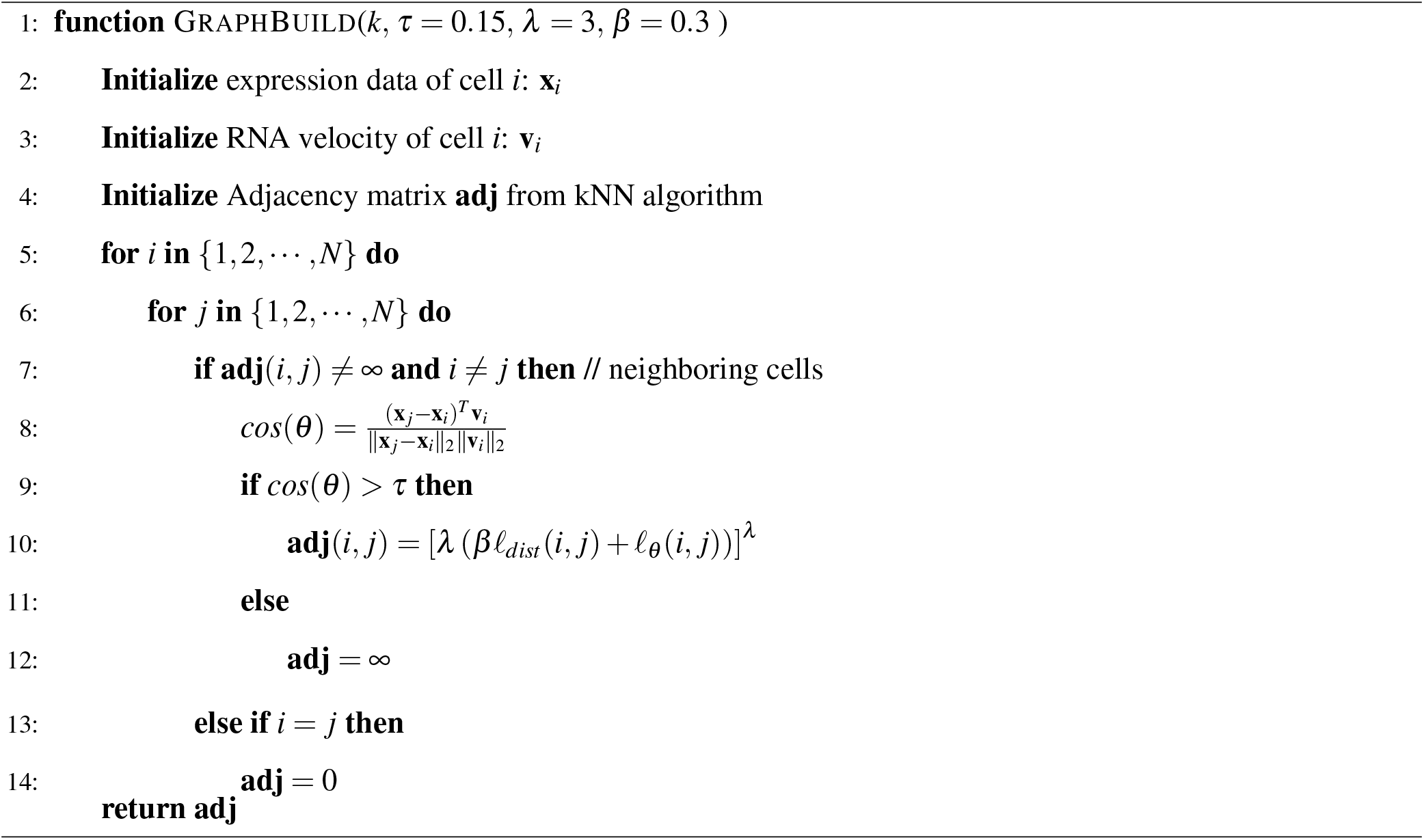
Build Graph

**Algorithm 2.**
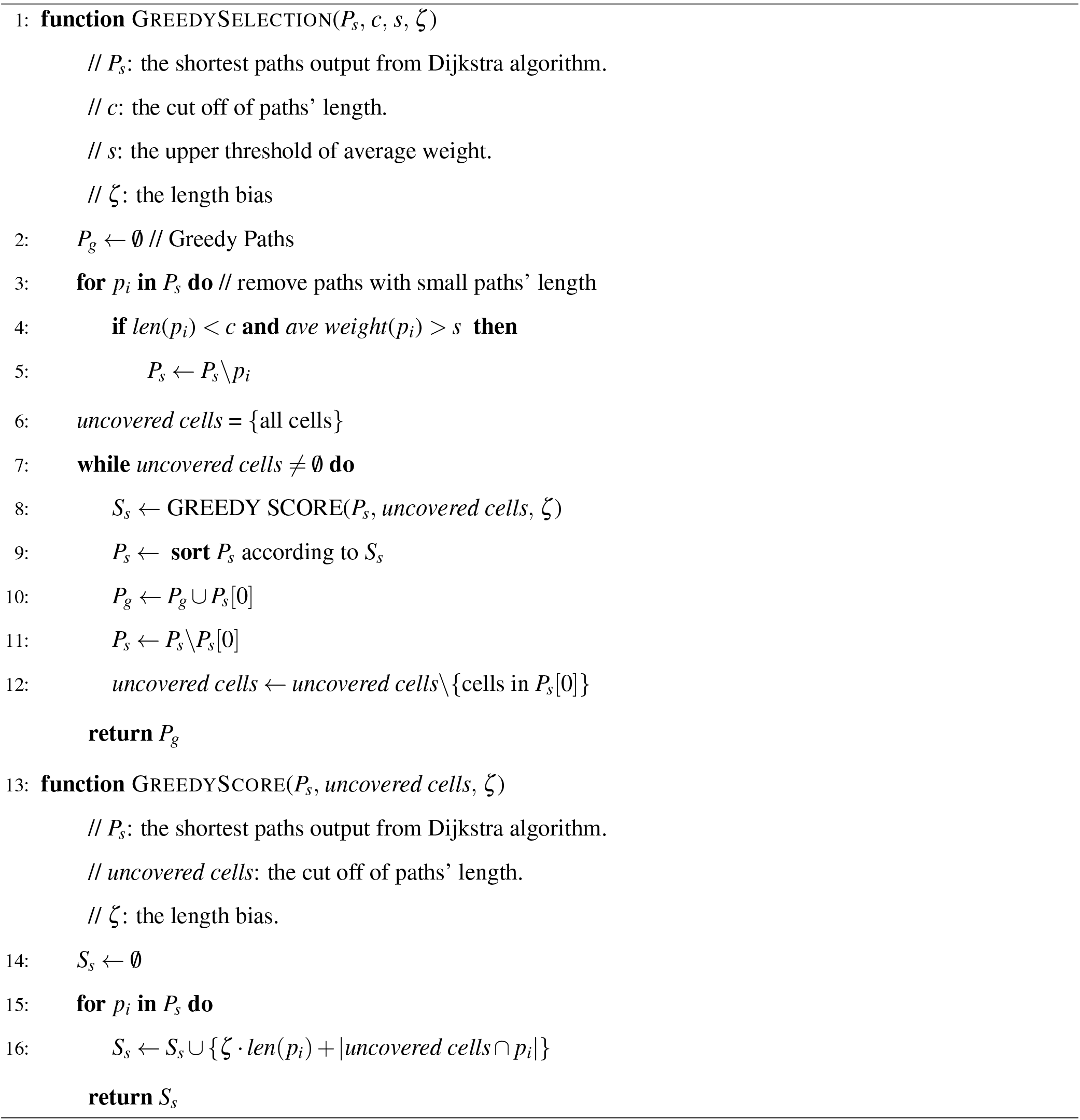
Greedy Paths Selection

